# Integrated assessment of fatty acid metabolism and cellular energy status using HILIC-MS/MS

**DOI:** 10.64898/2026.08.12.744242

**Authors:** Mariana Lopes, Kyle D. Roberts, Alexis E. Heath, Peder J. Lund

## Abstract

Acetyl-CoA and other acyl-CoA thioesters are critical intermediates in the metabolic reactions that cells rely on to produce energy and carry out biosynthesis. Therefore, the analysis of acyl-CoA provides valuable information about the metabolic activity of cells, especially when combined with stable isotope tracing. Acyl-CoA species are routinely monitored by reversed-phase liquid chromatography coupled to tandem mass spectrometry (RPLC-MS/MS). However, drastic differences in the hydrophobicity of short-chain versus long-chain acyl-CoA species have been challenging to accommodate with a single set of RPLC conditions. Here, we describe a convenient method based on hydrophilic interaction liquid chromatography (HILIC-MS/MS) for the concurrent detection of both short-chain and long-chain acyl-CoA and their corresponding acyl-carnitine species. Using this strategy, we tracked the metabolism of isotope-labeled fatty acids in multiple cell lines, which revealed differences in their propensities for fatty acid oxidation and the extent to which isotope incorporation into acyl-CoA mirrored that of acyl-carnitine. We also applied the HILIC-MS/MS workflow to the analysis of NADH and ATP, making it a useful technique for gauging cellular bioenergetics as reflected by the acetyl-CoA/CoA, NADH/NAD^+^, and ATP/ADP ratios. Altogether, this HILIC-MS/MS platform enables a streamlined analysis of acyl-CoA species and other key intermediates in cell metabolism.

## INTRODUCTION

All cells contend with the fundamental need to produce energy and synthesize the wide array of molecules that are essential for growth and survival. Acetyl-CoA fulfills a central role in these catabolic and anabolic processes by serving as a high-energy donor of carbon atoms in the form of acetyl groups.^1–3^ As the main driver of the TCA cycle in mitochondria, acetyl-CoA fuels the production of NADH to power the electron transport chain for ATP synthesis. Certain cell types, such as hepatocytes, may also preserve mitochondrial acetyl-CoA for future use via conversion to ketone bodies, which other tissues can metabolize back into acetyl-CoA. Outside of mitochondria, acetyl-CoA contributes to the synthesis of lipids and steroids, which are integral components of cell membranes and important building blocks for other molecules.

Fatty acid oxidation (FAO) is one of several pathways available for cells to produce acetyl-CoA.^4^ FAO begins with the uptake of free fatty acids from the extracellular environment through active or passive transport mechanisms.^5^ Within cells, fatty acids are transformed into acyl-CoA thioesters by acyl-CoA synthetases. These acyl-CoA species must then gain access to the mitochondrial matrix, which is the primary site of FAO. In contrast to short-chain acyl-CoA, long-chain acyl-CoA requires assistance from the carnitine shuttle system to do so.^6^ As part of this process, CPT1A/B, located on the outer mitochondrial membrane, transfers the acyl chain from acyl-CoA to carnitine, creating acyl-carnitine. Acyl-carnitine then transits to the inner mitochondrial membrane, where it is converted back into acyl-CoA by CPT2 and released into the mitochondrial matrix. Over successive rounds of FAO, the acyl chain is progressively shortened by two carbons, producing acetyl-CoA, NADH, and FADH_2_ as byproducts. This continues until the original acyl-CoA shrinks to a final length of two or three carbons, at which point the resulting acetyl-CoA from even-chain substrates or propionyl-CoA from odd-chain substrates cannot undergo further oxidation. Aside from FAO, the oxidation of pyruvate downstream of glycolysis also serves as a major source of acetyl-CoA.^1^ Under certain conditions, cells can make acetyl-CoA from ketone bodies, acetate, ethanol, and amino acids.

Owing to its significance in cell metabolism, acyl-CoA species are routine targets for analysis by LC-MS/MS. Standard workflows involve extraction with methanol or TCA, clean-up by SPE, and separation by reversed-phase chromatography at basic pH with detection by mass spectrometry.^7–14^ While these conditions have worked well for short-chain acyl-CoA like acetyl-CoA, the analysis of long-chain acyl-CoA has been more problematic due to peak tailing and overly strong retention.^15,16^ Different stationary phases (e.g, C8 instead of C18), phosphoric acid washes, and ion pairing reagents have all been examined as possible solutions to these issues.^15–25^ However, a modified approach with a less retentive column for long-chain acyl-CoA may not be retentive enough for their short-chain counterparts. Supplemental washing or conditioning steps have been shown to be effective at improving peak shape and eliminating carryover,^15,16^ but these prolong analysis time and reduce throughput. The use of ion pairing reagents can also enhance chromatographic performance,^19^ though this often leads to persistent contamination of the LC-MS system that may limit its ability to run other assay panels.

To date, few studies have explored HILIC as an alternative to reversed-phase chromatography for acyl-CoA analysis. This orthogonal mode of separation, which retains analytes based on polarity, was previously applied to profile hydroxycinnamoyl-CoA species from plants.^22^ Another study analyzed a broad range of acyl-CoA species using a dual-column system to perform sequential RPLC and HILIC runs.^15^ HILIC permitted separation of more polar species that were not adequately resolved by RPLC. Building on this prior work, here we describe the development of a streamlined, HILIC-based strategy for acyl-CoA analysis. The simplified workflow allows for the simultaneous detection of short-chain and long-chain acyl-CoA within a single run without a need for sample clean-up or special column treatments. The method is also suitable for acyl-carnitines, certain lipids, NADH, and ATP, making it broadly useful for obtaining a comprehensive assessment of fatty acid metabolism and the overall energy status of cells.

## MATERIALS AND METHODS

### Chemicals

Sodium butyrate, sodium butyrate-^13^C_4_ (#488380, 99% isotopic purity), sodium palmitate, and sodium palmitate-^13^C_16_ (#700258, 99% isotopic purity), acyl-CoA standards, acyl-carnitine standards, and the ADP standard were purchased from Millipore Sigma. Standards for NAD^+^, NADH, ATP, and AMP were purchased from Fisher Scientific. The PC 16:0/18:1 (1-palmitoyl-2-oleoyl-sn-glycero-3-phosphocholine) lipid standard was from Avanti Polar Lipids. Butyrate stocks were prepared in water at 400 mM. Palmitate stocks were prepared at 100 mM by heating to 60°C in 50% ethanol. Standards were dissolved in water, except for PC 16:0/18:1, which was dissolved in chloroform. For SILEC, light and heavy (^13^C_6_-^15^N_2_) calcium di-pantothenate (#P5155 and #705837, 98% isotopic purity) and light and heavy (2,6,7-^13^C_3_-[pyridyl-^15^N]) nicotinamide (#N0636 and #809799, 98% isotopic purity) were purchased from Millipore Sigma and dissolved in sterile water.

### Cell culture, isotope tracing, and SILEC

HCT116, Caco2, HT29, SW872, 93T449, and 293T cells were cultured in DMEM (ThermoFisher #11965092) with 10% FBS, 1× GlutaMAX, and 1× penicillin/streptomycin in a humidified incubator at 37°C and 5% CO_2_. For ^13^C isotope tracing, approximately 1 × 10^6^ cells were plated in 2 ml of complete medium in 6 well plates. After 1-2 days of growth, cell monolayers were rinsed in PBS and incubated in 1 ml of labeling medium, consisting of glucose-free DMEM (ThermoFisher #11966025) with 10% FBS, 1× GlutaMAX, and 1× penicillin/streptomycin supplemented with 800 μM unlabeled or ^13^C_4_-labeled butyrate or 200 μM unlabeled or ^13^C_16_-labeled palmitate. A higher concentration of butyrate was used to account for its lower carbon content as compared to palmitate, and ethanol was added to 0.1% to butyrate conditions to control for the ethanol present in the palmitate stocks. After 4 h of labeling, cells were washed with PBS and harvested by scraping in ice-cold 80% methanol on ice. For glucose starvation experiments, 3 × 10^5^ cells were plated in 1 ml of complete medium in 12 well plates. After 2 days of growth, cell monolayers were rinsed in PBS and then incubated overnight in complete medium or glucose-free DMEM as above. For SILEC, 293T cells were grown in a custom media formulation (Boca Scientific) based on DMEM (ThermoFisher #11965092) without glutamine, pantothenate, or nicotinamide. To produce isotope-labeled CoA species, media was supplemented with 32.8 μM light nicotinamide and 8.4 μM heavy calcium di-pantothenate. To produce isotope-labeled NAD^+^ and NADH, media was supplemented with 32.8 μM heavy nicotinamide and 8.4 μM light calcium di-pantothenate. In both cases, media was additionally supplemented with 10% dialyzed FBS, 1× GlutaMAX, and 1× penicillin/streptomycin. Cells were cultured in T-25 flasks at 1/20 split ratios for three passages to achieve a complete m+4 isotopic shift in CoA species and NAD^+^/NADH. After further expansion, SILEC cells were harvested by trypsinization, washed in PBS, and stored as frozen cell pellets for spike-in experiments.

### Metabolite extraction from cells

Cell pellets were resuspended in 500 μl ice-cold 80% methanol. For isotope-based quantification, SILEC-labeled cell pellets (pantothenate for CoA, nicotinamide for NAD^+^/NADH) were resuspended in 80% methanol and spiked into the extraction solvent. Standards were prepared alongside samples by adding unlabeled (light) standards to an equivalent volume of extraction solvent. Lysates from cell pellets or scrapings were cleared of precipitated protein and debris by centrifugation at 21,380 x g at 4°C for 10 mins. Supernatants (90% of initial extraction volume) were transferred to 96 deep-well plates and dried under a stream of nitrogen at 30°C (Biotage SPE Dry 96). For normalization of ATP levels, the protein precipitate was resolubilized in 1% SDS and 20 mM Tris pH 8 at 95°C for 20 mins and then assayed by BCA (Thermo). Metabolite extracts were reconstituted in a volume of 50% acetonitrile equal to 9% of the initial extraction volume (e.g., 45 μl from 500 μl extraction). The plate was sealed with a plastic film and then transferred to the LC autosampler.

### Metabolite extraction from mouse liver

Frozen mouse liver (30 mg) from our previous study^26^ was lyophilized and then processed into a fine powder with ZrO_2_ beads shaken in a Precellys Cryolys Evolution homogenizer (7200 rpm, 3 x 20 sec with 15 sec pauses, temperature 4°C). Ice-cold 80% methanol (500 μl) was then added, and the beads were shaken again (7200 rpm, 6 x 20 sec with 15 sec pauses, temperature 4°C). The recovered homogenate (400 μl) was precipitated with 4 volumes of 100% methanol and centrifuged at 21,380 x g at 4°C for 10 mins. The supernatant was then transferred to a deep well plate, dried under nitrogen, and reconstituted in 100 μl of 50% acetonitrile. The reconstituted sample was centrifuged at 21,380 x g at 4°C for 5 mins to remove insoluble debris, and the supernatant was transferred to an autosampler vial.

### LC-MS/MS for acyl-CoA, acyl-carnitine, phosphatidylcholine, and ATP

Metabolites were analyzed using an Agilent 1260 Infinity II LC system interfaced with a Thermo TSQ Altis triple quadrupole mass spectrometer. Resuspended samples were placed in an autosampler unit cooled to 4°C, and 1 μl of sample (2 μl for mouse liver) was injected for separation over an Agilent Poroshell 120 HILIC-Z column (3 x 100 mm, 2.7 μm) at a flow rate of 0.5 ml/min. Solvent A was 10 mM ammonium bicarbonate (pH 9.0) with 5 μM medronic acid in water. Solvent B was 90% acetonitrile with 10 mM ammonium bicarbonate (pH 9.0) and 5 μM medronic acid. The gradient consisted of 95% B for 0-1 min, 95-50% B for 1-7 mins, 50% B for 7-11 mins, 50-95% B for 11-12 mins, and 95% B for 12-15 mins. The column temperature was set to 40°C. An in-line 0.2 μm filter was placed downstream of the autosampler to protect the column. A divert valve directed LC flow to waste for the first and last minute of the gradient. General settings for electrospray ionization in positive mode were +3500 V, sheath gas 50, auxiliary gas 10, sweep gas 1, vaporizer temperature 350°C, and ion transfer tube temperature 325°C. SRM scans were performed with CID gas (argon) at 1.5 mTorr. Initial runs used a Q1 resolution of 0.7 and a Q3 resolution of 1.5. In the finalized methods, Q3 resolution was reduced to 1.2 for CoA analysis and 0.7 for isotope tracing and ATP analysis. Transition settings are outlined in Supplemental Tables S1-S6.

### LC-MS/MS for NAD^+^ and NADH

NAD^+^ and NADH were analyzed as above with the following modifications. Initial experiments used a gradient of 95% B for 0-3 mins, 95-50% B for 3-8 mins, 50% B for 8-13 mins, 50-95% B for 13-15 mins, and 95% B for 15-20 mins. Solvent B was 100% acetonitrile and solvent A was 10 mM ammonium bicarbonate (pH 9.0) with 5 μM medronic acid in water. The column temperature was set to 40°C. LC flow was diverted to waste for the first 4 mins and the last 1 min. Upon observing significant oxidation of NADH, an isocratic gradient of 65% B for 1.5 mins was tested with pure NAD^+^ and NADH standards. The column temperature was reduced to 27°C, and solvent A was adjusted to 10 mM ammonium bicarbonate (pH 9.5) and 5 μM medronic acid. No divert valve was used. The final method for analyzing NADH in cell extracts, which is amenable to other nicotinamide-related metabolites,^27^ used the same mobile phases as for CoA, carnitine, and ATP analysis (A = 10 mM ammonium bicarbonate [pH 9.0] + 5 μM medronic acid; B = 90% acetonitrile + 10 mM ammonium bicarbonate [pH 9.0] + 5 μM medronic acid). To account for the lower acetonitrile content of solvent B, the gradient was modified to 70% B for 0-1 mins, 70-50% B for 1-2 mins, 50% B for 2-5 mins, 50-70% B for 5-6 mins, and 70% B for 6-10 mins. The column temperature was kept at 27°C, and the first 0.5 min and the last minute of the gradient was diverted to waste. Dwell time was 80 ms, and Q1 and Q3 resolutions were 0.4 and 1.2, respectively.

### Data analysis

MS raw files were imported into Skyline to calculate peak areas for each transition. This data was exported for further analysis in Excel, Prism, and R. All chromatograms were plotted in R using retention time and intensity values from Skyline. For isotope tracing experiments, the peak areas of isotopologues were converted into percent molar enrichment using the online FluxFix calculator^28^ and peak areas from controls treated with unlabeled fatty acids. For isotope-based quantification of acyl-CoA and NAD^+^/NADH, the peak areas of the light (L, m+0) and heavy (H, m+4) precursors were calculated as the sum of the two (acyl-CoA) or three (NAD^+^/NADH) contributing fragment ions. The endogenous light precursors were normalized by the area of the heavy spike-in (L/H ratio). The L/H ratios of unknown samples were then used to calculate concentrations based on linear regression of the internal calibration curve, constructed from L/H ratios of known standards. In the case of ATP with an external calibration curve, metabolite amounts were normalized to the amount of protein recovered in the precipitate. Statistical testing was performed in Prism.

## RESULTS

### Development of a HILIC-MS/MS method for acyl-CoA and acyl-carnitines

To monitor fatty acid oxidation and cellular energetics, we aimed to develop an LC-MS workflow capable of analyzing both long-chain and short-chain acyl-CoA and acyl-carnitine species. Most approaches for these targets have relied on reversed-phase chromatography for upstream separation. However, species with shorter acyl chains are less hydrophobic and are generally not retained well by reversed-phase columns. The use of ion pairing reagents can promote retention of more polar molecules, but this may suppress electrospray ionization and limit compatibility of the LC-MS system with other methods since these reagents are difficult to remove. In contrast to the poor retention of molecules with short-chain acyl groups, those with longer acyl chains may bind too strongly to reversed-phase columns. In our initial work, we noted that the peaks for palmitoyl-CoA and palmitoyl-carnitine were much weaker than for other species under the reversed-phase conditions that we tested (**SUP. FIG. S1A-B**). We also observed minimal retention of carnitine species with acyl chains numbering less than four carbons (**SUP. FIG. S1B**).

In light of these drawbacks, we explored HILIC as an alternative separation method. This strategy effectively resolved pure standards of short-chain and long-chain acyl-CoA and acyl-carnitines as well as their unconjugated forms (**FIG. 1A-J**). As expected for HILIC-based separations, species with longer acyl chains and lower polarity eluted earlier in the gradient. We also applied this method to analyze acyl-CoA and acyl-carnitines extracted from murine liver tissue as a representative source of biological material (**FIG. 1K-L**). Consistent with prior work on acyl-CoA and acyl-carnitines in positive mode ESI, we detected common fragment ions at 428 *m/z* and 85 *m/z*, respectively.^12,29^ However, the dominant fragment for all acyl-CoA species, except palmitoyl-CoA (**FIG. 1D**), arose from the neutral loss of ADP, which is well-established.^12^ This variable fragment (e.g., 303 *m/z* for acetyl-CoA) corresponds to the pantothenate backbone linked to the acyl group, meaning that its mass reflects the length of the acyl chain. For all acyl-carnitines, the signature fragment ion at 85 *m/z* (C_4_H_5_O_2_ ^+^) is most dominant, which corresponds to the carnitine backbone after loss of the trimethylamine and acyl moieties.^29^ However, as for acyl-CoA species, we also detected a variable fragment ion arising from the neutral loss of trimethylamine. Since it retains the acyl group, the mass of this fragment (e.g., 145 *m/z* for acetyl-carnitine) is indicative of the acyl chain length.

**FIGURE 1.**
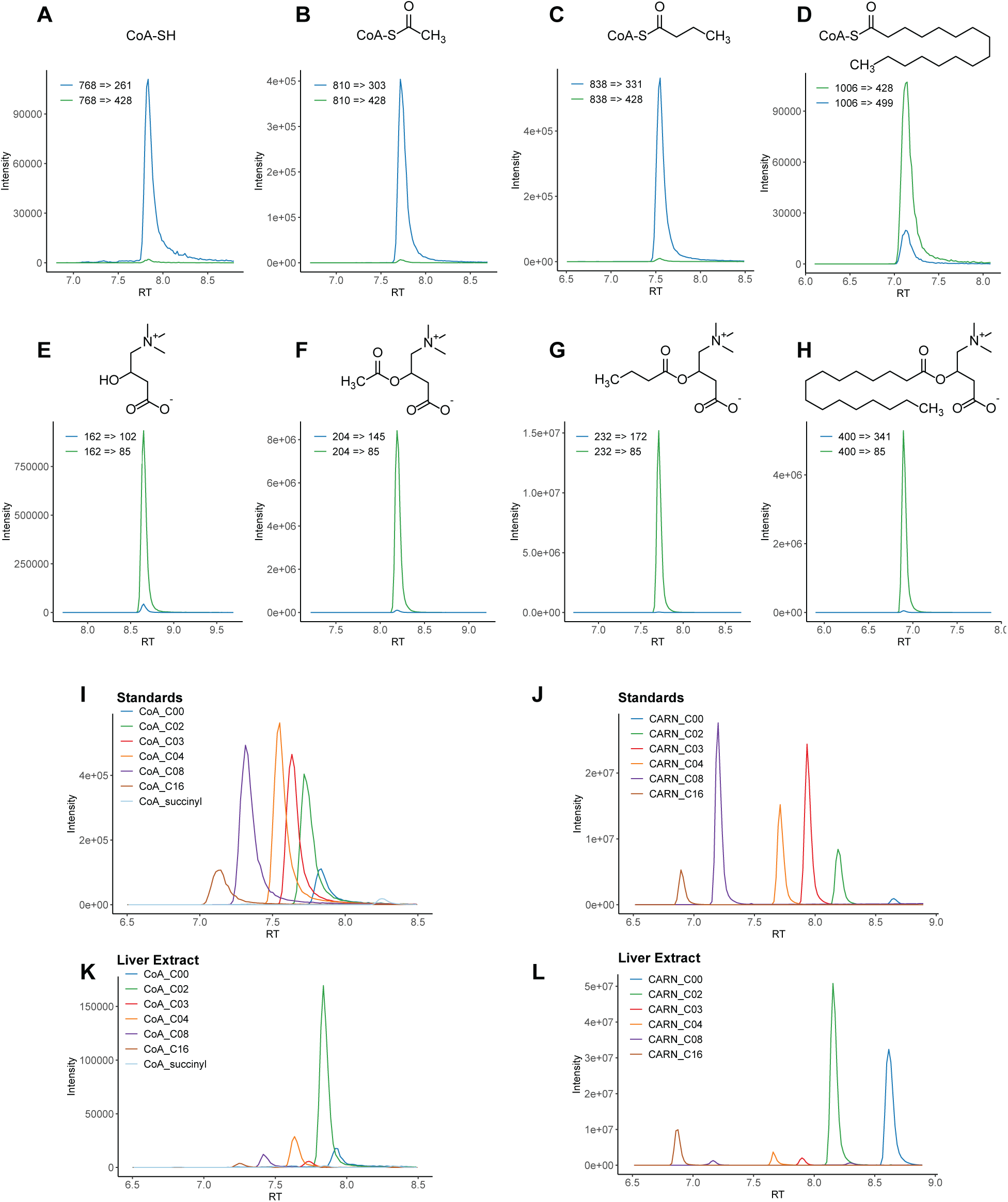
Development of a HILIC-MS/MS method for acyl-CoA and acyl-carnitines. A standard mix containing 1.5 μM of each acyl-CoA and 0.5 μM of each acyl-carnitine was resolved by HILIC for detection by MS/MS with a QqQ instrument in positive mode. Representative traces of two transitions each (common and variable) for (**A**) CoA, (**B**) acetyl-CoA, (**C**) butyryl-CoA, (**D**) palmitoyl-CoA, (**E**) carnitine, (**F**) acetyl-carnitine, (**G**) butyryl-carnitine, and (**H**) palmitoyl-carnitine. Overlaid traces of the most intense transition for (**I**) acyl-CoA and (**J**) acyl-carnitine species of varying chain lengths. Overlaid traces for (**K**) acyl-CoA and (**L**) acyl-carnitine species extracted from mouse liver tissue.

### Tracing the oxidation of fatty acids into acetyl-CoA

As an application of this new method, we compared the metabolic processing of two fatty acid substrates in different cell lines by stable isotope tracing. For these experiments, we chose butyrate and palmitate as prototypical short-chain and long-chain fatty acids, respectively. Butyrate is a major byproduct of the gut microbiota that is considered to be the main energy source for epithelial cells in the colon.^30^ Palmitate is a highly abundant saturated fatty acid that can be produced through de novo lipogenesis or absorbed from the diet.^31,32^ After treating cells with ^13^C-labeled butyrate or palmitate, we analyzed acyl-CoA and acyl-carnitines for isotope incorporation. To accomplish this, we performed SRM scans targeting the isotopologues of precursor ions and their corresponding fragment ions that carry ^13^C-labeled acyl groups derived from butyrate or palmitate. As expected, we detected a strong signal from the M+16 isotopologue of palmitoyl-CoA in HT29 cells treated with ^13^C-labeled but not unlabeled palmitate, indicating efficient uptake and metabolic activation of this long-chain fatty acid into its CoA thioester (**FIG. 2A**). We also detected a significant increase in the M+2 isotopologue of acetyl-CoA, which is evidence of palmitate being catabolized into two-carbon units through fatty acid oxidation (**FIG. 2B**).

Acetyl-CoA can enter the TCA cycle to drive the generation of ATP through oxidative phosphorylation. Alternatively, acetyl-CoA can contribute to lipid synthesis. We found that the HILIC-MS/MS method was also capable of detecting phospholipids, such as phosphatidylcholine 16:0/18:1 (**FIG. 2C**), which allowed us to track the catabolic and anabolic metabolism of fatty acids simultaneously. In addition to oxidizing palmitate into acetyl-CoA, we noted that HT29 cells incorporated this fatty acid into PC 16:0/18:1, as evident by a more intense M+16 isotopologue after treatment with ^13^C-palmitate (**FIG. 2C**). With the method adapted for isotope tracing analysis, we proceeded to compare ^13^C labeling patterns from palmitate and butyrate across different cell lines (**FIG. 2G-J**).

**FIGURE 2.**
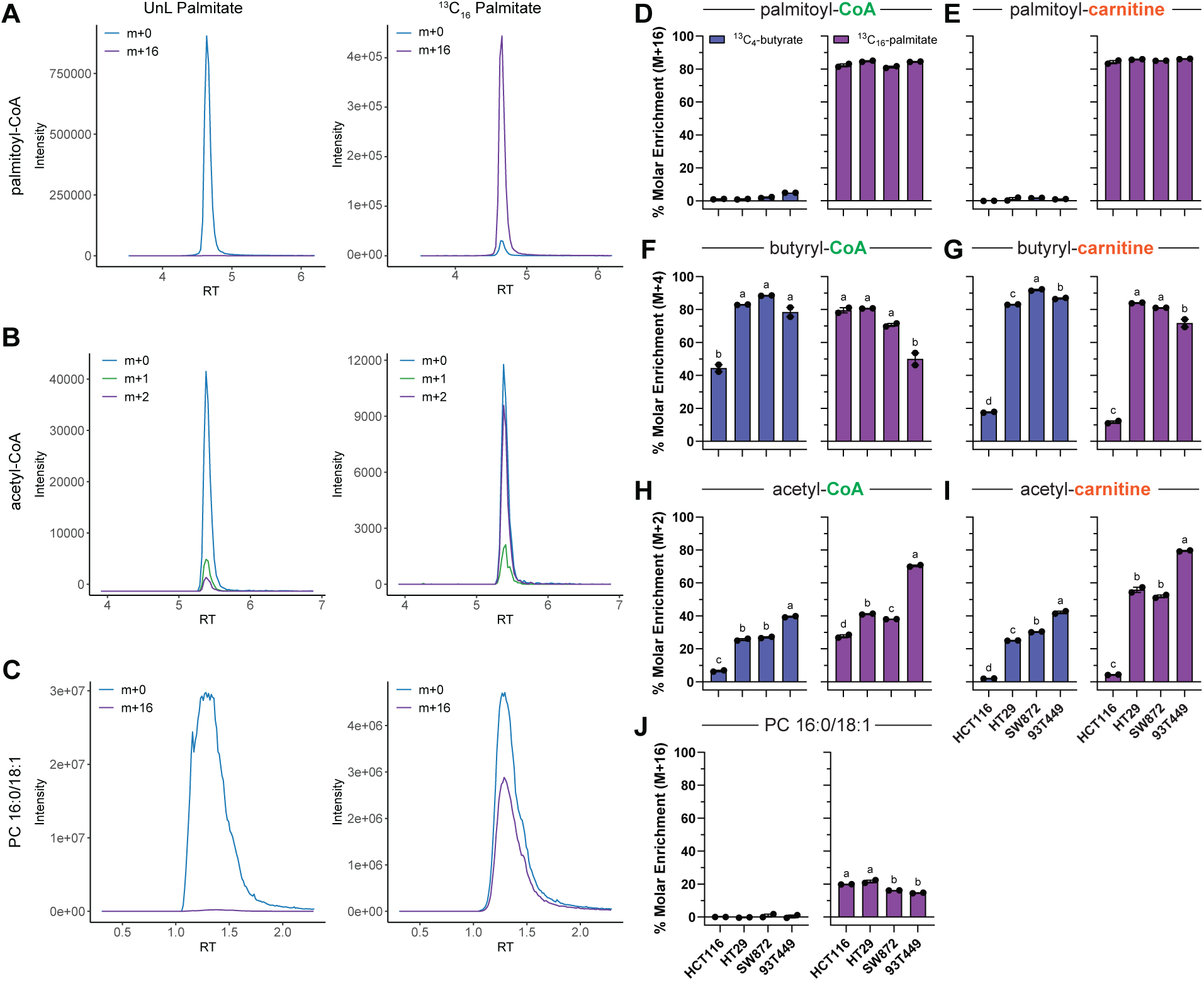
Tracing the oxidation of fatty acids into acetyl-CoA. Cell lines were incubated with 800 μM unlabeled or ^13^C-labeled butyrate or 200 μM unlabeled or ^13^C-labeled palmitate for 4 hrs. Traces of selected isotopologues (*m+0*, *m+1*, *m+2*, *m+16*) of (**A**) palmitoyl-CoA, (**B**) acetyl-CoA, or (**C**) phosphatidylcholine 16:0/18:1 from HT29 cells treated with unlabeled or ^13^C-labeled palmitate. Percent molar enrichment of fully labeled (**D**) palmitoyl-CoA, (**E**) palmitoyl-carnitine, (**F**) butyryl-CoA, (**G**) butyryl-carnitine, (**H**) acetyl-CoA, (**I**) acetyl-carnitine, (**J**) phosphatidylcholine 16:0/18:1 after treatment of cell lines with ^13^C_4_-butyrate or ^13^C_16_-palmitate. Mean ± sem, n = 2. Bars with different letters have statistically significant differences in their means based on one-way ANOVA with Tukey post-hoc tests comparing the labeling across different cell lines from a given fatty acid.

Palmitoyl-CoA and palmitoyl-carnitine incorporated the ^13^C label from palmitate to a similar extent in all cells, suggesting no major differences in the initial transport and processing of this fatty acid (**FIG. 2D-E**). In contrast, the pools of labeled butyryl-CoA from butyrate, but not palmitate, were significantly lower in HCT116 cells (**FIG. 2F**). These cells also showed substantially less isotope incorporation into butyryl-carnitine. This was surprising because butyryl-carnitine was robustly labeled in other cell lines that attained similar levels of butyryl-CoA labeling as HCT116 cells (**FIG. 2F-G**). Both butyrate and palmitate underwent oxidation into acetyl-CoA in all cell lines (**FIG. 2H**). The molar enrichment of heavy acetyl-CoA in 93T449 cells reached 70% from palmitate and nearly 40% from butyrate. These values dropped to 30% and 7% in HCT116 cells, which again showed only minor amounts of the corresponding ^13^C-labeled acyl-carnitine species (**FIG. 2H-I**). Palmitate contributed to phosphatidylcholine synthesis in all cell lines (**FIG. 2J**). In contrast, we did not observe notable enrichment of any isotopologues from PC 16:0/18:1 after ^13^C-butyrate treatment, indicating that this short-chain fatty acid did not contribute meaningfully to lipid synthesis.

### Isotope-based quantification of acetyl-CoA/CoA ratio

Acetyl-CoA is a key intermediate in energy metabolism. The acetyl group arises from the catabolic processing of numerous carbon sources, including fatty acids, glucose-derived pyruvate, acetate, ketone bodies, ethanol, and amino acids.^1^ Thus, the ratio of acetylated versus free CoA represents a useful measure of cellular energy status. As another application of the method, we sought to assess cellular energetics by quantifying levels of acyl-CoA. Quantitative LC-MS assays often rely on isotope-labeled internal standards for enhanced accuracy and precision. While heavy standards for acyl-CoA species are available from commercial suppliers, we generated our own standards through SILEC labeling.^8,33^ This approach entails growing cells in heavy pantothenate, which forms the backbone of CoA. As a result, all acyl-CoA species acquire isotopic labeling that introduces a mass shift of 4 Da. These heavy SILEC cells can then be spiked into samples and synthetic standards of known concentration, which only contain light acyl-CoA (M+0). Peak areas of the light species (L) are then normalized to those of the heavy spike-in (H), yielding a L/H ratio that is more resistant to technical variation and easily converted into an absolute quantity based on a calibration curve.

To assess our ability to measure changes in the acetyl-CoA/CoA ratio, we cultured cells in the absence of glucose, which should reduce the production of acetyl-CoA from pyruvate. As expected, the signal from endogenous acetyl-CoA (M+0) relative to the SILEC spike-in (M+4) decreased in response to glucose starvation (**FIG. 3A**). Using a calibration curve, we calculated the concentrations of acetyl-CoA and CoA in cell extracts and found a significant drop in the ratio of acetylated versus free CoA, attesting to the validity of our approach (**FIG. 3B-C**).

**FIGURE 3.**
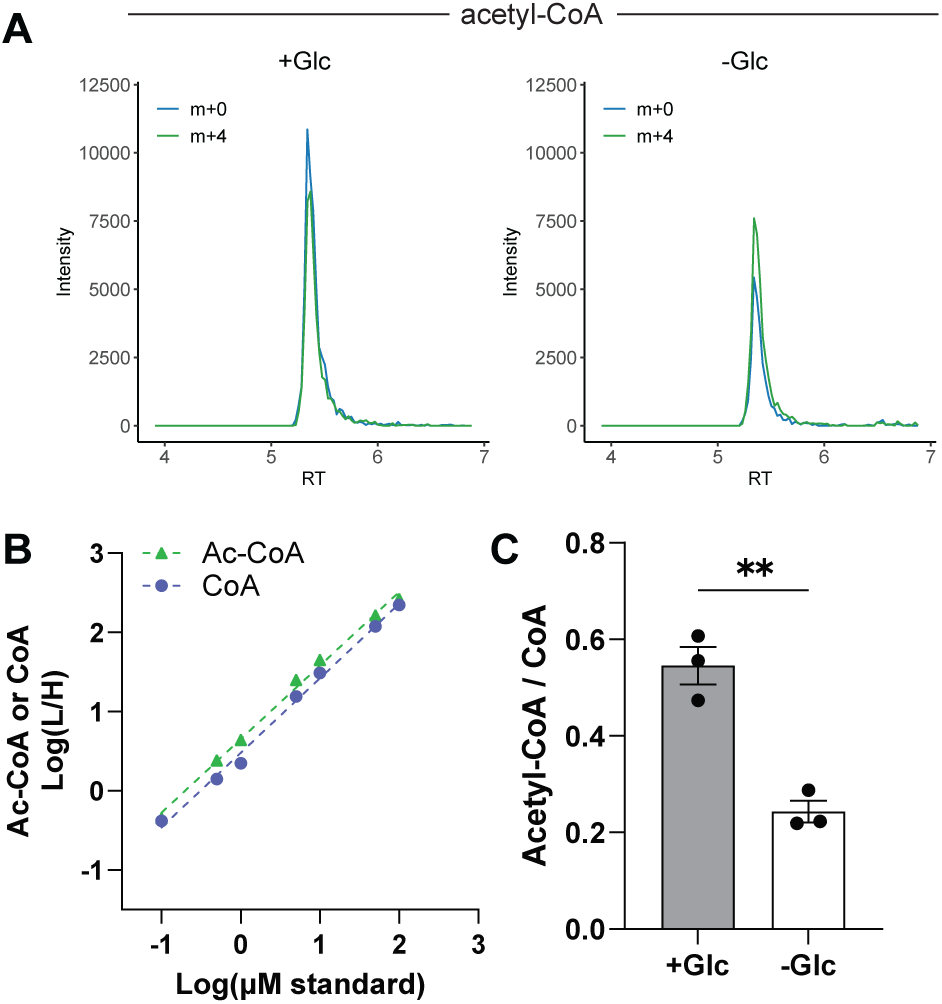
Isotope-based quantification of acetyl-CoA/CoA ratio. Caco2 cells were starved of glucose overnight and then lysed in extraction solvent spiked with pantothenate-labeled SILEC cells. (**A**) Traces of acetyl-CoA from Caco2 cells (*m+0*) under normal (*+Glc*) and starved (*-Glc*) conditions overlaid with the acetyl-CoA signal (*m+4*) from the heavy SILEC spike-in. (**B**) Internal calibration curves comparing concentrations of light standards to their heavy-normalized peak areas (*L/H ratio*). R^2^ = 0.9945 for CoA and 0.9954 for acetyl-CoA. (**C**) Ratio of acetyl-CoA to CoA in normal and starved Caco2 cells. Mean ± sem, n = 3. ** p < 0.01 by unpaired t-test.

### Isotope-based quantification of NADH/NAD^+^ ratio

Like the acetyl-CoA/CoA ratio, the balance of oxidized NAD^+^ to reduced NADH also provides information about cellular energetics and redox state. NADH is generated during glycolysis, fatty acid oxidation, and the TCA cycle. One of its main functions is to carry electrons to the electron transport chain to power ATP synthesis. Seeking to establish an integrated LC-MS panel for a more comprehensive assessment of cellular energy status, we examined whether the HILIC method for acyl-CoA and acyl-carnitines was also applicable to NADH. Although we obtained well-resolved peaks for both NAD^+^ and NADH, a significant NAD^+^ signal appeared in the NADH standard (**FIG. 4A**). Conversely, no NADH signal was observed in the NAD^+^ standard. Direct infusion of the NADH standard confirmed its purity, and injection of increasingly concentrated NADH standards correlated with stronger NAD^+^ signals. These findings pointed to artifactual conversion of NADH to NAD^+^ during the HILIC gradient rather than a contaminated sample or carryover of residual NAD^+^ from prior injections. Spurious oxidation of NADPH during HILIC has been mentioned in the literature.^34^

To counter this issue, which would otherwise produce an erroneous NADH/NAD^+^ ratio, we tuned the HILIC conditions to a lower percentage of mobile phase B to minimize the retention time of NADH while maintaining adequate separation from NAD^+^ (**FIG. 4B**). Although this adjustment did not completely eliminate the spurious NAD^+^ peak, it significantly attenuated its intensity to under 10% of the signal from the true NAD^+^ standard. In this analysis, we also detected a minor NAD^+^ peak at the same retention time as NADH, likely from in-source oxidation. To control for these remaining artifacts, we again turned to a SILEC strategy in which we grew cells in heavy nicotinamide to introduce an isotopic shift of 4 Da in NAD^+^ and NADH, as previously reported.^35^ We reasoned that any artificial increase or decrease in the NAD^+^ or NADH peak, respectively, would happen equivalently to a spike-in of heavy SILEC cells, preserving the original L/H ratios. These L/H ratios would then facilitate accurate measurement of NADH and NAD^+^.

To validate this workflow, we analyzed cells that had been deprived of glucose as a model of energy deficiency with less NADH production. As shown in **FIG. 4C-D**, we used the peak areas of light standards spiked with heavy-labeled SILEC cells to construct calibration curves for NAD^+^ and NADH based on L/H ratios. As expected, we found the NADH/NAD^+^ ratio to be significantly lower in two cell lines after glucose starvation (**FIG. 4E-F**). Notably, the NADH/NAD^+^ ratio that we calculated for HCT116 cells under standard glucose conditions (0.149 ± 0.016, mean ± sd) is in line with a previously published value (0.172 ± 0.009).^36^

**FIGURE 4.**
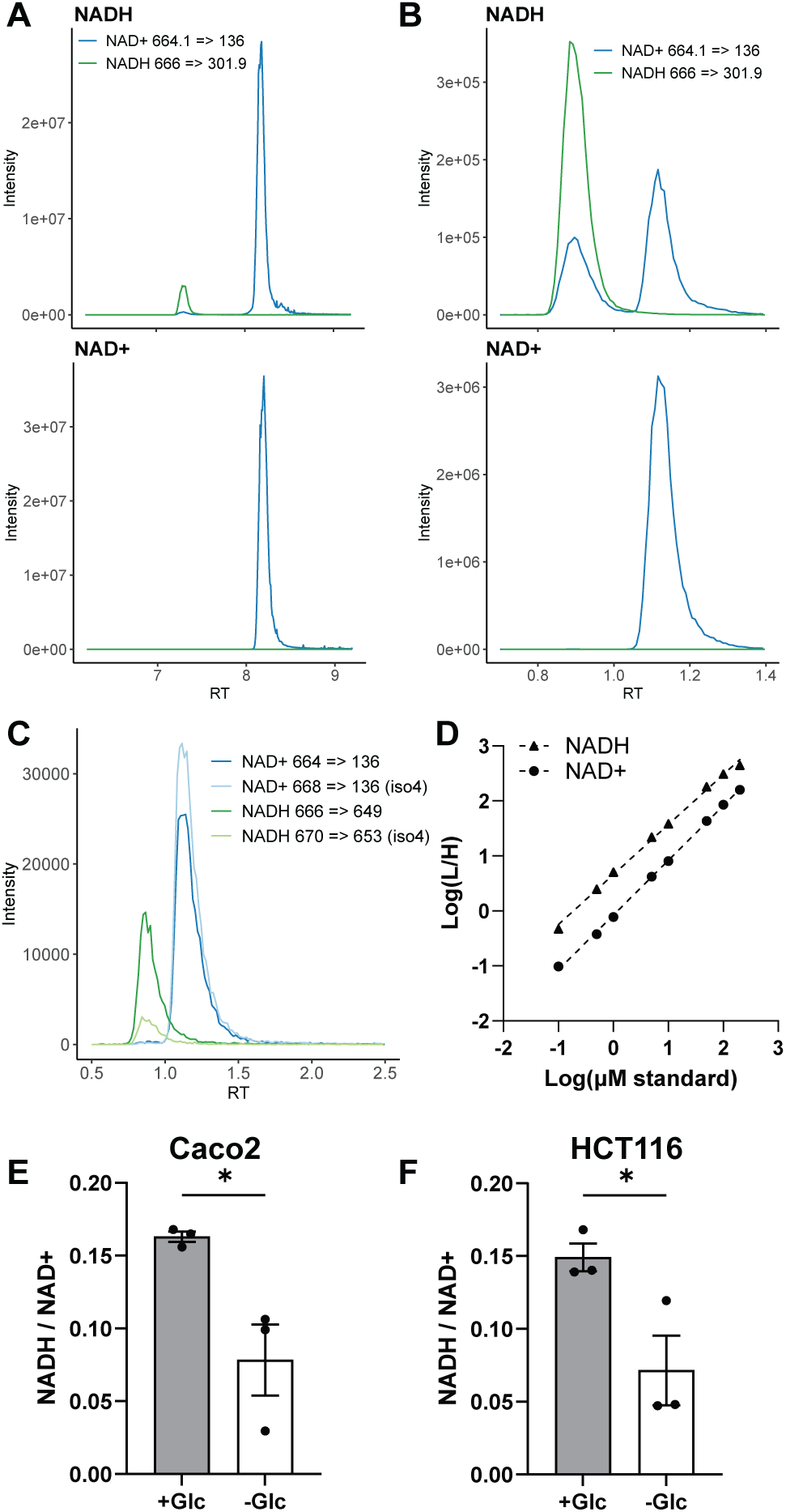
Isotope-based quantification of NADH/NAD^+^ ratio. (**A**) NADH and NAD^+^ signals in 1 μM standards of NADH (*top*) or NAD^+^ (*bottom*) analyzed with a 20 min HILIC gradient, showing significant oxidation of NADH to NAD^+^. (**B**) NADH and NAD^+^ signals in 10 μM standards analyzed with a modified gradient to minimize retention time and limit artificial oxidation. (**C**) Representative traces of light NADH and NAD^+^ in a 1 μM standard mix overlaid with NADH and NAD^+^ signals (*iso4*) from nicotinamide-labeled SILEC cells. (**D**) Internal calibration curves comparing concentrations of light standards to their heavy-normalized peak areas (*L/H ratio*). R^2^ = 0.9969 for NADH and 0.9991 for NAD^+^. (**E**) Caco2 and (**F**) HCT116 cells were starved of glucose overnight and then lysed in extraction solvent spiked with nicotinamide-labeled SILEC cells. Ratio of reduced NADH to oxidized NAD^+^ in normal (*+Glc*) and starved (*-Glc*) cells. Mean ± sem, n = 3. * p < 0.05 by unpaired t-test.

### Assessment of ATP/ADP ratio

Acetyl-CoA and NADH ultimately support the synthesis of ATP from ADP. The ATP/ADP ratio is perhaps the most fundamental indicator of the overall energy status of cells. Therefore, we explored the possibility of including ATP, ADP, and AMP as targets in our HILIC method. We obtained well-defined peaks for these molecules from synthetic standards as well as cell extracts (**FIG. 5A-B**). As expected, the presence of more phosphate groups correlated with longer retention times. Using a calibration curve built from external standards (**FIG. 5C**), we quantified the levels of adenosine nucleotides in cells starved of glucose, which we predicted would suppress ATP production. While these starved cells had a modest but statistically significant decrease in total ATP levels, they maintained the same ATP/ADP ratio as controls (**FIG. 5D-E**). We also calculated the adenylate energy charge (AEC),^37^ which approaches a value of one as more AMP and ADP are converted into ATP. Similar to control cells, glucose-starved cells had an AEC of roughly 0.9 (**SUP. FIG. S2A**), which falls within the normal range of 0.75-0.9.^38^

**FIGURE 5.**
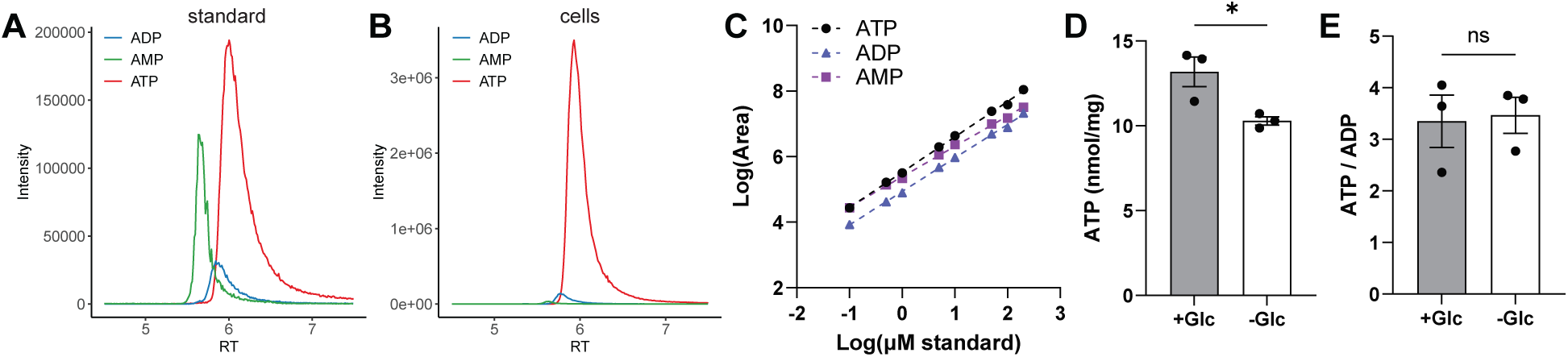
Assessment of ATP/ADP ratio. Traces of ATP, ADP, and AMP from (**A**) a standard mix containing 10 μM of each and (**B**) an extract from HCT116 cells. (**C**) External calibration curves comparing concentrations of standards to their peak areas. R^2^ = 0.9987 for ATP, 0.9987 for ADP, and 0.9985 for AMP. Measurement of (**D**) ATP levels and (**E**) ATP/ADP ratio in normal (*+Glc*) and glucose-starved (*-Glc*) HCT116 cells. Mean ± sem, n = 3. * p < 0.05 by unpaired t-test.

We were surprised that glucose deprivation did not have a significant impact on the ATP/ADP ratio or AEC in light of the observed decline in acetyl-CoA and NADH (**FIG. 3C, 4E-F**). One possibility is that cells reduced their demand for ATP by arresting proliferation and other energy-consuming processes. In support of this scenario, we recovered less total protein from glucose-starved cells (**SUP. FIG. S2B**). Cells may have also catabolized glutamine or other amino acids, which were not restricted like glucose. Glutaminolysis produces α-ketoglutarate, which can then enter the TCA cycle.^39,40^ However, fueling the TCA cycle with glutamine instead of glucose-derived pyruvate generates less NADH overall, which matches the trends in our data. Upstream of the TCA, the conversion of glucose into two molecules of acetyl-CoA through pyruvate decarboxylation results in four equivalents of NADH. Together, these two molecules of acetyl-CoA then yield six equivalents of NADH through the TCA cycle. In contrast, metabolism of glutamine into α-KG by glutaminase and glutamate dehydrogenase only produces one molecule of NADH. Entering the TCA cycle downstream of acetyl-CoA and citrate, this α-KG can then produce two or three equivalents of NADH, depending on whether it meets with a source of acetyl-CoA to continue the cycle or exits as malate. Likewise, acetyl-CoA production from glutamine is typically much lower than from glucose-derived pyruvate because it requires energy to operate the TCA cycle in reverse or the diversion of malate.^41^ Nonetheless, glutamine metabolism may have been sufficient to maintain the levels of ATP, but not NADH and acetyl-CoA, in glucose-starved cells on a temporary basis. A longer period of glucose deprivation, potentially with glutamine restriction, may lead to a more dramatic effect on ATP levels in HCT116 cells.

## CONCLUSION

In summary, we have developed a reliable and easily implemented HILIC-MS/MS platform with broad utility for the rapid analysis of key intermediates in cellular energy metabolism, including acyl-CoA, acyl-carnitine, NADH/NAD^+^, and ATP/ADP/AMP. This approach complements existing methods that have largely employed reversed-phase chromatography, assisted by ion pairing reagents in some cases.^7,25^ A major strength of our HILIC-based strategy is that it captures both short-chain and long-chain acyl-CoA and acyl-carnitines in a single 15-minute run with a single set of mobile phases, overcoming the challenge associated with their varying degrees of hydrophobicity. Upstream sample preparation is similarly convenient and straightforward, involving a simple methanol-based extraction without a requirement for additional clean-up steps.

Although the same HILIC setup is compatible with NADH analysis, a minor drawback is the need to run a separate, less retentive gradient because of the artificial oxidation that occurs with longer retention times. However, any use of SILEC-based internal standards would already necessitate separate processing workflows. For instance, a spike-in of nicotinamide-labeled SILEC cells with heavy NADH/NAD^+^ would also contain unlabeled acyl-CoA and ATP, which would overlap with the acyl-CoA and ATP signals from the sample being interrogated. Nonetheless, all runs can still take place within the same queue since they do not require any hardware changes. Future work could explore a multiplexed SILEC strategy to introduce isotopic shifts on all analytes of interest (e.g., double labeling with pantothenate and nicotinamide), creating a universal spike-in standard. Even though each sample would still need to be injected twice to analyze NADH on a different gradient than acyl-CoA and ATP, this would at least allow for a single processing workflow.

As a biological application, we combined this new HILIC-MS/MS method with stable isotope tracing to compare the metabolism of two ^13^C-labeled fatty acid substrates with different chain lengths. This analysis revealed variations in the propensity of different cell lines for fatty acid oxidation and a striking disconnect between acyl-CoA and acyl-carnitine labeling in HCT116 cells. A lack of carnitine O-acetyltransferase activity, which mediates the reversible transfer of acyl groups between CoA and carnitine,^1^ could potentially account for the latter finding. In another application, we used isotope-based quantification to measure acetyl-CoA/CoA and NADH/NAD^+^ ratios as readouts of cellular energy status, both of which decline during acute nutrient starvation. Notably, we were able to conduct all analyses in an economical manner with a triple quadrupole instrument. Altogether, this HILIC-MS/MS platform provides a convenient and unified approach for the analysis of central intermediates in fatty acid oxidation and energy metabolism.

## SUPPORTING INFORMATION

- Transition lists; RPLC analysis of acyl-CoA and acyl-carnitine; and adenylate energy charge and protein levels after glucose starvation.

## ACKNOWLEDGEMENTS

PJL acknowledges support from institutional start-up funds, the Cleveland DDRCC (P30DK097948), the Case Comprehensive Cancer Center in partnership with the American Cancer Society (IRG-22-148-27-IRG), and the Willard A. Bernbaum Cystic Fibrosis Research Center.

## CONTRIBUTIONS

**ML**: conceptualization, methodology, investigation, formal analysis, and writing (review and editing). **KDR** and **AEH**: methodology, investigation, formal analysis, and writing (review and editing). **PJL**: project conceptualization, methodology, investigation, formal analysis, visualization, writing (original draft and editing), supervision, and funding acquisition.

## Supporting Information

**Supplemental Table S1.**
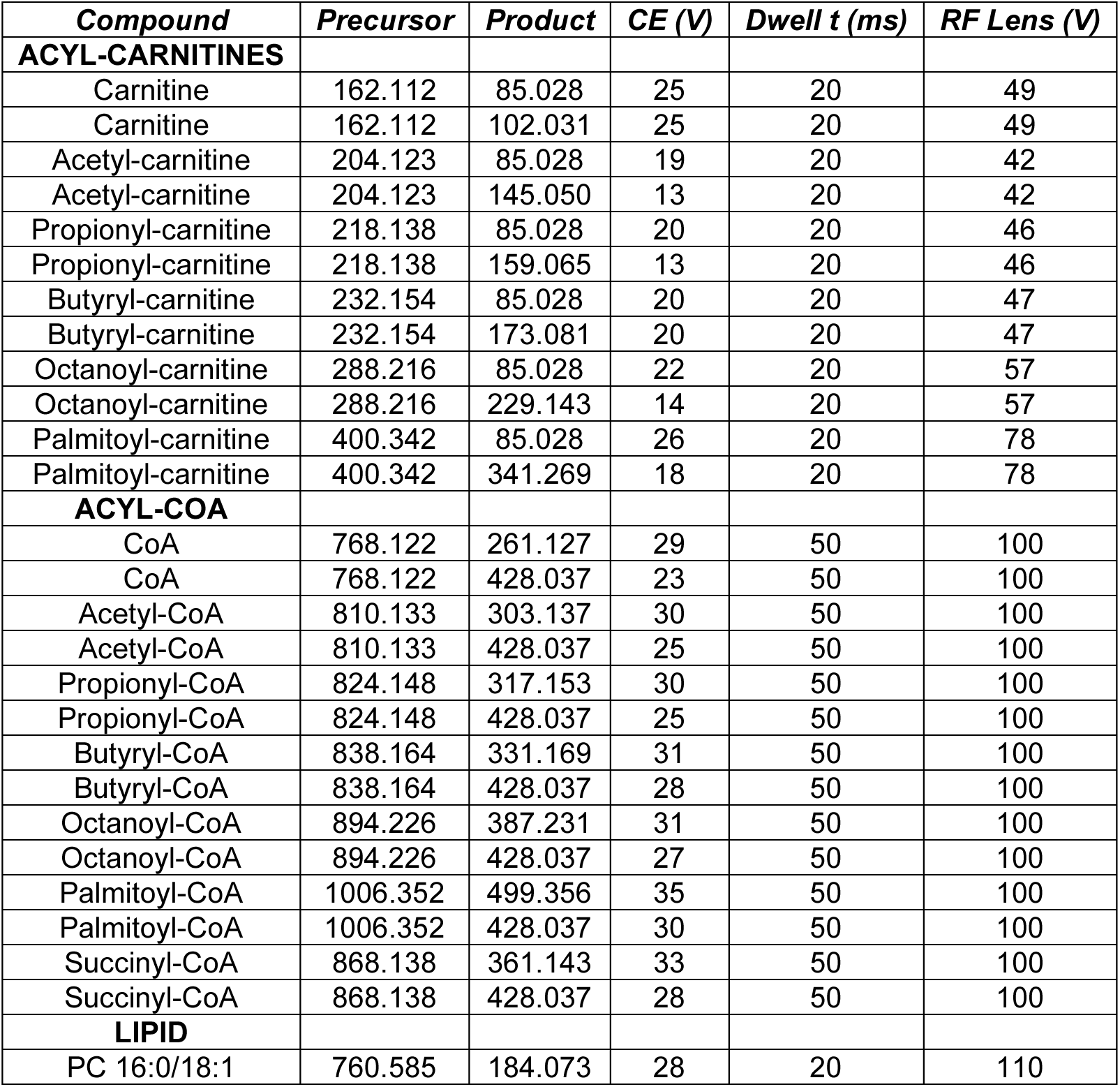
List of transitions for the general analysis of acyl-carnitine, acyl-CoA, and PC 16:0/18:1.

| ***Compound*** | ***Precursor*** | ***Product*** | ***CE (V)*** | ***Dwell t (ms)*** | ***RF Lens (V)*** |
| --- | --- | --- | --- | --- | --- |
| **ACYL-CARNITINES** |  |  |  |  |  |
| Carnitine | 162.112 | 85.028 | 25 | 20 | 49 |
| Carnitine | 162.112 | 102.031 | 25 | 20 | 49 |
| Acetyl-carnitine | 204.123 | 85.028 | 19 | 20 | 42 |
| Acetyl-carnitine | 204.123 | 145.050 | 13 | 20 | 42 |
| Propionyl-carnitine | 218.138 | 85.028 | 20 | 20 | 46 |
| Propionyl-carnitine | 218.138 | 159.065 | 13 | 20 | 46 |
| Butyryl-carnitine | 232.154 | 85.028 | 20 | 20 | 47 |
| Butyryl-carnitine | 232.154 | 173.081 | 20 | 20 | 47 |
| Octanoyl-carnitine | 288.216 | 85.028 | 22 | 20 | 57 |
| Octanoyl-carnitine | 288.216 | 229.143 | 14 | 20 | 57 |
| Palmitoyl-carnitine | 400.342 | 85.028 | 26 | 20 | 78 |
| Palmitoyl-carnitine | 400.342 | 341.269 | 18 | 20 | 78 |
| **ACYL-COA** |  |  |  |  |  |
| CoA | 768.122 | 261.127 | 29 | 50 | 100 |
| CoA | 768.122 | 428.037 | 23 | 50 | 100 |
| Acetyl-CoA | 810.133 | 303.137 | 30 | 50 | 100 |
| Acetyl-CoA | 810.133 | 428.037 | 25 | 50 | 100 |
| Propionyl-CoA | 824.148 | 317.153 | 30 | 50 | 100 |
| Propionyl-CoA | 824.148 | 428.037 | 25 | 50 | 100 |
| Butyryl-CoA | 838.164 | 331.169 | 31 | 50 | 100 |
| Butyryl-CoA | 838.164 | 428.037 | 28 | 50 | 100 |
| Octanoyl-CoA | 894.226 | 387.231 | 31 | 50 | 100 |
| Octanoyl-CoA | 894.226 | 428.037 | 27 | 50 | 100 |
| Palmitoyl-CoA | 1006.352 | 499.356 | 35 | 50 | 100 |
| Palmitoyl-CoA | 1006.352 | 428.037 | 30 | 50 | 100 |
| Succinyl-CoA | 868.138 | 361.143 | 33 | 50 | 100 |
| Succinyl-CoA | 868.138 | 428.037 | 28 | 50 | 100 |
| **LIPID** |  |  |  |  |  |
| PC 16:0/18:1 | 760.585 | 184.073 | 28 | 20 | 110 |

**Supplemental Table S2.**
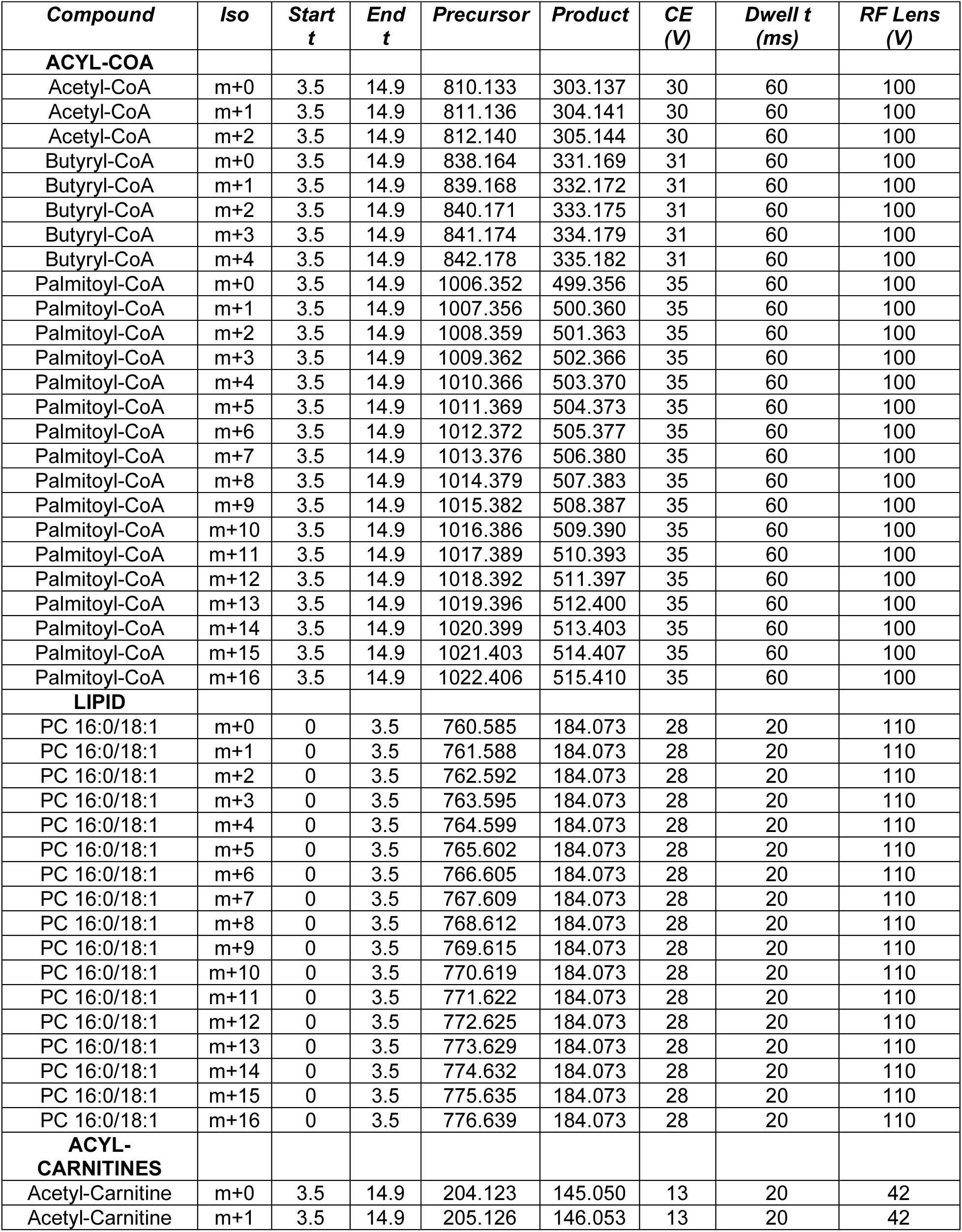
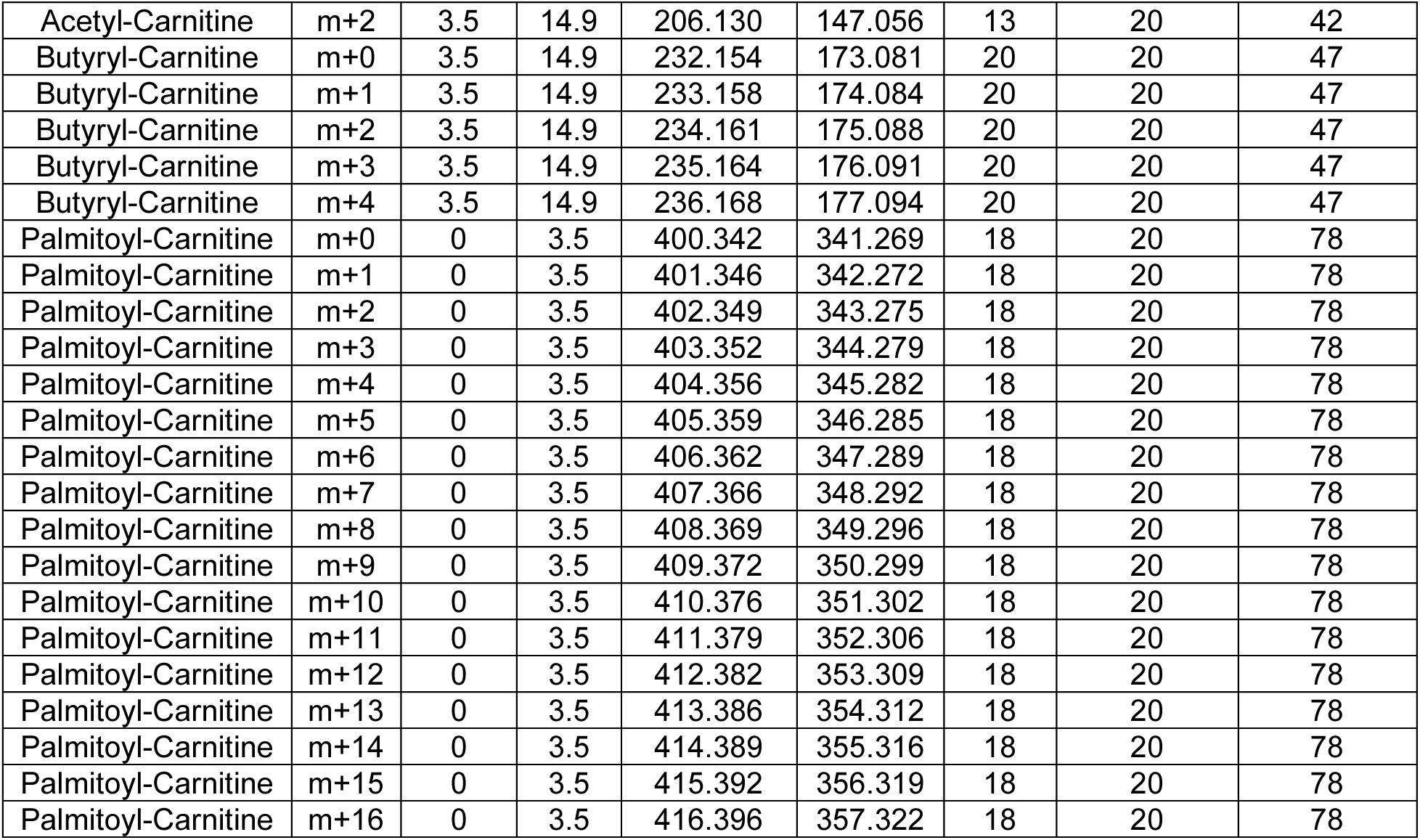
List of transitions for isotope tracing analysis targeting acyl-CoA, acyl-carnitine, and PC 16:0/18:1.

| *Compound* | *Iso* | *Start*<br>*t* | *End*<br>*t* | *Precursor* | *Product* | *CE*<br>*(V)* | *Dwell t*<br>*(ms)* | *RF Lens*<br>*(V)* |
| --- | --- | --- | --- | --- | --- | --- | --- | --- |
| **ACYL-COA** |  |  |  |  |  |  |  |  |
| Acetyl-CoA | m+0 | 3.5 | 14.9 | 810.133 | 303.137 | 30 | 60 | 100 |
| Acetyl-CoA | m+1 | 3.5 | 14.9 | 811.136 | 304.141 | 30 | 60 | 100 |
| Acetyl-CoA | m+2 | 3.5 | 14.9 | 812.140 | 305.144 | 30 | 60 | 100 |
| Butyryl-CoA | m+0 | 3.5 | 14.9 | 838.164 | 331.169 | 31 | 60 | 100 |
| Butyryl-CoA | m+1 | 3.5 | 14.9 | 839.168 | 332.172 | 31 | 60 | 100 |
| Butyryl-CoA | m+2 | 3.5 | 14.9 | 840.171 | 333.175 | 31 | 60 | 100 |
| Butyryl-CoA | m+3 | 3.5 | 14.9 | 841.174 | 334.179 | 31 | 60 | 100 |
| Butyryl-CoA | m+4 | 3.5 | 14.9 | 842.178 | 335.182 | 31 | 60 | 100 |
| Palmitoyl-CoA | m+0 | 3.5 | 14.9 | 1006.352 | 499.356 | 35 | 60 | 100 |
| Palmitoyl-CoA | m+1 | 3.5 | 14.9 | 1007.356 | 500.360 | 35 | 60 | 100 |
| Palmitoyl-CoA | m+2 | 3.5 | 14.9 | 1008.359 | 501.363 | 35 | 60 | 100 |
| Palmitoyl-CoA | m+3 | 3.5 | 14.9 | 1009.362 | 502.366 | 35 | 60 | 100 |
| Palmitoyl-CoA | m+4 | 3.5 | 14.9 | 1010.366 | 503.370 | 35 | 60 | 100 |
| Palmitoyl-CoA | m+5 | 3.5 | 14.9 | 1011.369 | 504.373 | 35 | 60 | 100 |
| Palmitoyl-CoA | m+6 | 3.5 | 14.9 | 1012.372 | 505.377 | 35 | 60 | 100 |
| Palmitoyl-CoA | m+7 | 3.5 | 14.9 | 1013.376 | 506.380 | 35 | 60 | 100 |
| Palmitoyl-CoA | m+8 | 3.5 | 14.9 | 1014.379 | 507.383 | 35 | 60 | 100 |
| Palmitoyl-CoA | m+9 | 3.5 | 14.9 | 1015.382 | 508.387 | 35 | 60 | 100 |
| Palmitoyl-CoA | m+10 | 3.5 | 14.9 | 1016.386 | 509.390 | 35 | 60 | 100 |
| Palmitoyl-CoA | m+11 | 3.5 | 14.9 | 1017.389 | 510.393 | 35 | 60 | 100 |
| Palmitoyl-CoA | m+12 | 3.5 | 14.9 | 1018.392 | 511.397 | 35 | 60 | 100 |
| Palmitoyl-CoA | m+13 | 3.5 | 14.9 | 1019.396 | 512.400 | 35 | 60 | 100 |
| Palmitoyl-CoA | m+14 | 3.5 | 14.9 | 1020.399 | 513.403 | 35 | 60 | 100 |
| Palmitoyl-CoA | m+15 | 3.5 | 14.9 | 1021.403 | 514.407 | 35 | 60 | 100 |
| Palmitoyl-CoA | m+16 | 3.5 | 14.9 | 1022.406 | 515.410 | 35 | 60 | 100 |
| **LIPID** |  |  |  |  |  |  |  |  |
| PC 16:0/18:1 | m+0 | 0 | 3.5 | 760.585 | 184.073 | 28 | 20 | 110 |
| PC 16:0/18:1 | m+1 | 0 | 3.5 | 761.588 | 184.073 | 28 | 20 | 110 |
| PC 16:0/18:1 | m+2 | 0 | 3.5 | 762.592 | 184.073 | 28 | 20 | 110 |
| PC 16:0/18:1 | m+3 | 0 | 3.5 | 763.595 | 184.073 | 28 | 20 | 110 |
| PC 16:0/18:1 | m+4 | 0 | 3.5 | 764.599 | 184.073 | 28 | 20 | 110 |
| PC 16:0/18:1 | m+5 | 0 | 3.5 | 765.602 | 184.073 | 28 | 20 | 110 |
| PC 16:0/18:1 | m+6 | 0 | 3.5 | 766.605 | 184.073 | 28 | 20 | 110 |
| PC 16:0/18:1 | m+7 | 0 | 3.5 | 767.609 | 184.073 | 28 | 20 | 110 |
| PC 16:0/18:1 | m+8 | 0 | 3.5 | 768.612 | 184.073 | 28 | 20 | 110 |
| PC 16:0/18:1 | m+9 | 0 | 3.5 | 769.615 | 184.073 | 28 | 20 | 110 |
| PC 16:0/18:1 | m+10 | 0 | 3.5 | 770.619 | 184.073 | 28 | 20 | 110 |
| PC 16:0/18:1 | m+11 | 0 | 3.5 | 771.622 | 184.073 | 28 | 20 | 110 |
| PC 16:0/18:1 | m+12 | 0 | 3.5 | 772.625 | 184.073 | 28 | 20 | 110 |
| PC 16:0/18:1 | m+13 | 0 | 3.5 | 773.629 | 184.073 | 28 | 20 | 110 |
| PC 16:0/18:1 | m+14 | 0 | 3.5 | 774.632 | 184.073 | 28 | 20 | 110 |
| PC 16:0/18:1 | m+15 | 0 | 3.5 | 775.635 | 184.073 | 28 | 20 | 110 |
| PC 16:0/18:1 | m+16 | 0 | 3.5 | 776.639 | 184.073 | 28 | 20 | 110 |
| **ACYL-CARNITINES** |  |  |  |  |  |  |  |  |
| Acetyl-Carnitine | m+0 | 3.5 | 14.9 | 204.123 | 145.050 | 13 | 20 | 42 |
| Acetyl-Carnitine | m+1 | 3.5 | 14.9 | 205.126 | 146.053 | 13 | 20 | 42 |
| Acetyl-Carnitine | m+2 | 3.5 | 14.9 | 206.130 | 147.056 | 13 | 20 | 42 |
| Butyryl-Carnitine | m+0 | 3.5 | 14.9 | 232.154 | 173.081 | 20 | 20 | 47 |
| Butyryl-Carnitine | m+1 | 3.5 | 14.9 | 233.158 | 174.084 | 20 | 20 | 47 |
| Butyryl-Carnitine | m+2 | 3.5 | 14.9 | 234.161 | 175.088 | 20 | 20 | 47 |
| Butyryl-Carnitine | m+3 | 3.5 | 14.9 | 235.164 | 176.091 | 20 | 20 | 47 |
| Butyryl-Carnitine | m+4 | 3.5 | 14.9 | 236.168 | 177.094 | 20 | 20 | 47 |
| Palmitoyl-Carnitine | m+0 | 0 | 3.5 | 400.342 | 341.269 | 18 | 20 | 78 |
| Palmitoyl-Carnitine | m+1 | 0 | 3.5 | 401.346 | 342.272 | 18 | 20 | 78 |
| Palmitoyl-Carnitine | m+2 | 0 | 3.5 | 402.349 | 343.275 | 18 | 20 | 78 |
| Palmitoyl-Carnitine | m+3 | 0 | 3.5 | 403.352 | 344.279 | 18 | 20 | 78 |
| Palmitoyl-Carnitine | m+4 | 0 | 3.5 | 404.356 | 345.282 | 18 | 20 | 78 |
| Palmitoyl-Carnitine | m+5 | 0 | 3.5 | 405.359 | 346.285 | 18 | 20 | 78 |
| Palmitoyl-Carnitine | m+6 | 0 | 3.5 | 406.362 | 347.289 | 18 | 20 | 78 |
| Palmitoyl-Carnitine | m+7 | 0 | 3.5 | 407.366 | 348.292 | 18 | 20 | 78 |
| Palmitoyl-Carnitine | m+8 | 0 | 3.5 | 408.369 | 349.296 | 18 | 20 | 78 |
| Palmitoyl-Carnitine | m+9 | 0 | 3.5 | 409.372 | 350.299 | 18 | 20 | 78 |
| Palmitoyl-Carnitine | m+10 | 0 | 3.5 | 410.376 | 351.302 | 18 | 20 | 78 |
| Palmitoyl-Carnitine | m+11 | 0 | 3.5 | 411.379 | 352.306 | 18 | 20 | 78 |
| Palmitoyl-Carnitine | m+12 | 0 | 3.5 | 412.382 | 353.309 | 18 | 20 | 78 |
| Palmitoyl-Carnitine | m+13 | 0 | 3.5 | 413.386 | 354.312 | 18 | 20 | 78 |
| Palmitoyl-Carnitine | m+14 | 0 | 3.5 | 414.389 | 355.316 | 18 | 20 | 78 |
| Palmitoyl-Carnitine | m+15 | 0 | 3.5 | 415.392 | 356.319 | 18 | 20 | 78 |
| Palmitoyl-Carnitine | m+16 | 0 | 3.5 | 416.396 | 357.322 | 18 | 20 | 78 |

**Supplemental Table S3.**
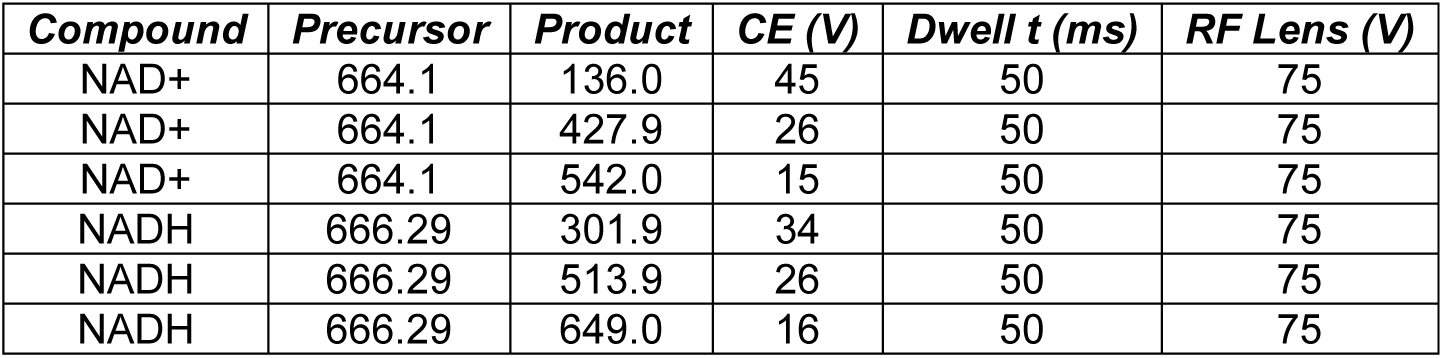
List of transitions for NADH/NAD^+^ analysis.

| *Compound* | *Precursor* | *Product* | *CE (V)* | *Dwell t (ms)* | *RF Lens (V)* |
| --- | --- | --- | --- | --- | --- |
| NAD<sup>+</sup> | 664.1 | 136.0 | 45 | 50 | 75 |
| NAD<sup>+</sup> | 664.1 | 427.9 | 26 | 50 | 75 |
| NAD<sup>+</sup> | 664.1 | 542.0 | 15 | 50 | 75 |
| NADH | 666.29 | 301.9 | 34 | 50 | 75 |
| NADH | 666.29 | 513.9 | 26 | 50 | 75 |
| NADH | 666.29 | 649.0 | 16 | 50 | 75 |

**Supplemental Table S4.** List of transitions for isotope-based quantification of acetyl-CoA/CoA ratio.

| *Compound* | *Iso* | *Precursor* | *Product* | *CE (V)* | *Dwell t (ms)* | *RF Lens (V)* |
| --- | --- | --- | --- | --- | --- | --- |
| CoA | m+0 | 768.1 | 261.1 | 29 | 50 | 100 |
| CoA | m+0 | 768.1 | 428.0 | 23 | 50 | 100 |
| CoA | m+4 | 772.1 | 265.1 | 29 | 50 | 100 |
| CoA | m+4 | 772.1 | 428.0 | 23 | 50 | 100 |
| Acetyl-CoA | m+0 | 810.1 | 303.1 | 30 | 50 | 100 |
| Acetyl-CoA | m+0 | 810.1 | 428.0 | 25 | 50 | 100 |
| Acetyl-CoA | m+4 | 814.1 | 307.1 | 30 | 50 | 100 |
| Acetyl-CoA | m+4 | 814.1 | 428.0 | 25 | 50 | 100 |
| Propionyl-CoA | m+0 | 824.1 | 317.2 | 30 | 50 | 100 |
| Propionyl-CoA | m+0 | 824.1 | 428.0 | 25 | 50 | 100 |
| Propionyl-CoA | m+4 | 828.2 | 321.2 | 30 | 50 | 100 |
| Propionyl-CoA | m+4 | 828.2 | 428.0 | 25 | 50 | 100 |
| Butyryl-CoA | m+0 | 838.2 | 331.2 | 31 | 50 | 100 |
| Butyryl-CoA | m+0 | 838.2 | 428.0 | 26 | 50 | 100 |
| Butyryl-CoA | m+4 | 842.2 | 335.2 | 31 | 50 | 100 |
| Butyryl-CoA | m+4 | 842.2 | 428.0 | 26 | 50 | 100 |
| Succinyl-CoA | m+0 | 868.1 | 361.1 | 33 | 50 | 100 |
| Succinyl-CoA | m+0 | 868.1 | 428.0 | 28 | 50 | 100 |
| Succinyl-CoA | m+4 | 872.1 | 365.1 | 33 | 50 | 100 |
| Succinyl-CoA | m+4 | 872.1 | 428.0 | 28 | 50 | 100 |
| Octanoyl-CoA | m+0 | 894.2 | 387.2 | 31 | 50 | 100 |
| Octanoyl-CoA | m+0 | 894.2 | 428.0 | 27 | 50 | 100 |
| Octanoyl-CoA | m+4 | 898.2 | 391.2 | 31 | 50 | 100 |
| Octanoyl-CoA | m+4 | 898.2 | 428.0 | 27 | 50 | 100 |
| Palmitoyl-CoA | m+0 | 1006.4 | 499.4 | 35 | 50 | 100 |
| Palmitoyl-CoA | m+0 | 1006.4 | 428.0 | 30 | 50 | 100 |
| Palmitoyl-CoA | m+4 | 1010.4 | 503.4 | 35 | 50 | 100 |
| Palmitoyl-CoA | m+4 | 1010.4 | 428.0 | 30 | 50 | 100 |

**Supplemental Table S5.** List of transitions for isotope-based quantification of NADH/NAD^+^ ratio.

| *Compound* | *Iso* | *Precursor* | *Product* | *CE (V)* | *Dwell t (ms)* | *RF Lens (V)* |
| --- | --- | --- | --- | --- | --- | --- |
| NAD<sup>+</sup> | m+0 | 664.1 | 136.0 | 45 | 80 | 75 |
| NAD<sup>+</sup> | m+0 | 664.1 | 427.9 | 26 | 80 | 75 |
| NAD<sup>+</sup> | m+0 | 664.1 | 542.0 | 15 | 80 | 75 |
| NAD<sup>+</sup> | m+4 | 668.1 | 136.0 | 45 | 80 | 75 |
| NAD<sup>+</sup> | m+4 | 668.1 | 427.9 | 26 | 80 | 75 |
| NAD<sup>+</sup> | m+4 | 668.1 | 542.0 | 15 | 80 | 75 |
| NADH | m+0 | 666.1 | 301.9 | 34 | 80 | 75 |
| NADH | m+0 | 666.1 | 513.9 | 26 | 80 | 75 |
| NADH | m+0 | 666.1 | 649.0 | 16 | 80 | 75 |
| NADH | m+4 | 670.1 | 305.9 | 34 | 80 | 75 |
| NADH | m+4 | 670.1 | 517.9 | 26 | 80 | 75 |
| NADH | m+4 | 670.1 | 653.0 | 16 | 80 | 75 |

**Supplemental Table S6.** List of transitions for ATP, ADP, and AMP.

| ***Compound*** | ***Precursor*** | ***Product*** | ***CE (V)*** | ***Dwell t (ms)*** | ***RF Lens (V)*** |
| --- | --- | --- | --- | --- | --- |
| ATP | 508 | 136.0 | 31 | 50 | 75 |
| ATP | 508 | 348.0 | 16 | 50 | 75 |
| ATP | 508 | 409.9 | 17 | 50 | 75 |
| ADP | 428 | 118.9 | 55 | 50 | 67 |
| ADP | 428 | 136.0 | 24 | 50 | 67 |
| ADP | 428 | 348.0 | 17 | 50 | 67 |
| AMP | 348 | 97.0 | 30 | 50 | 64 |
| AMP | 348 | 119.0 | 55 | 50 | 64 |
| AMP | 348 | 136.0 | 20 | 50 | 64 |

**Supplemental Figure S1.**
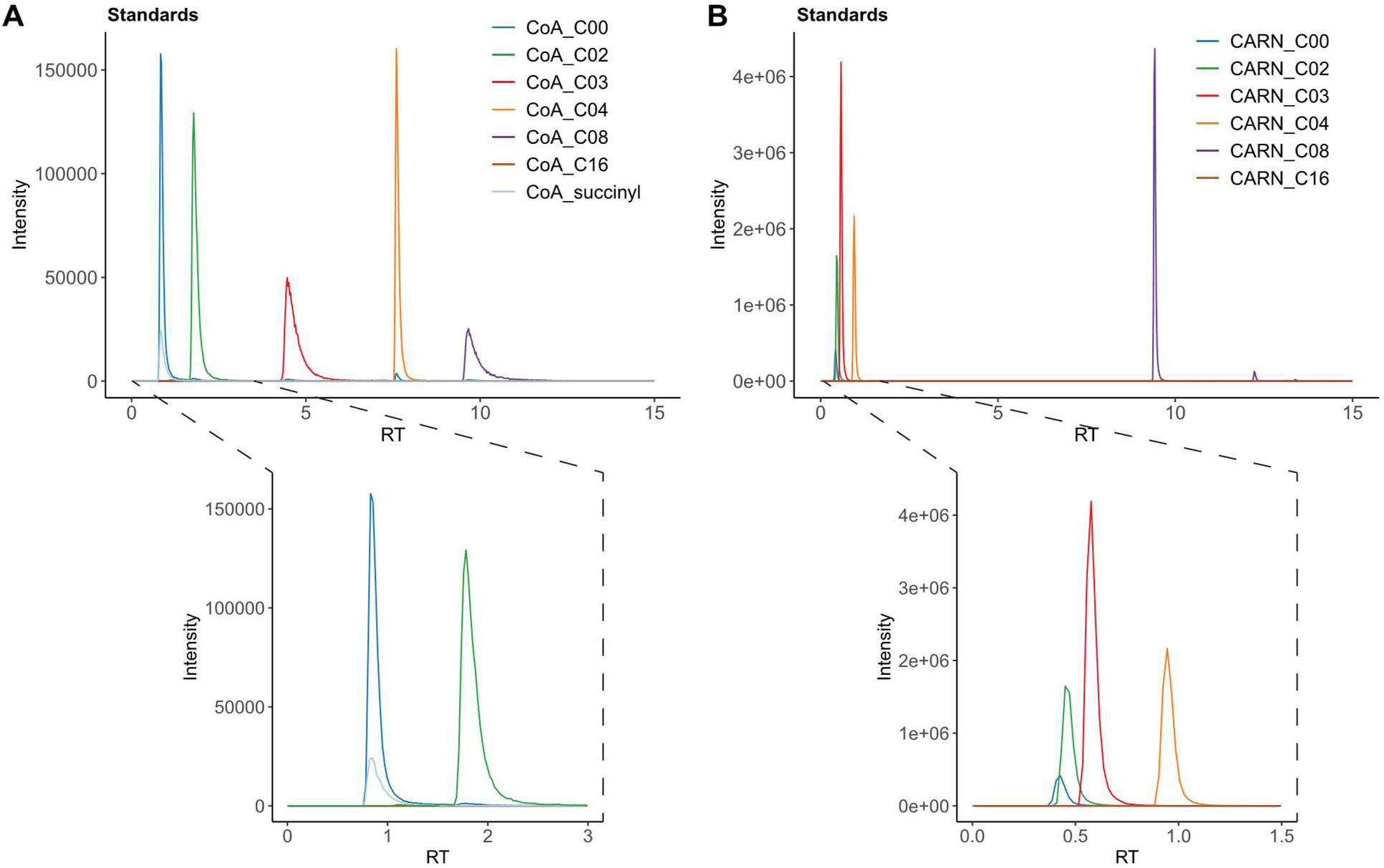
RPLC analysis of acyl-CoA and acyl-carnitine. A standard mixture containing (**A**) 5 µM of each acyl-CoA or (**B**) 0.5 µM of each acyl-carnitine was resolved on a Waters Atlantis BEH C18 AX column (2.1 x 50 mm, 1.7 µm) at a flow rate of 0.5 ml/min and a temperature of 40°C. The injection volume was 1 µl. Solvent A was 10 mM ammonium bicarbonate pH 9.0 in water, and solvent B was acetonitrile. The gradient consisted of 5% B from 0-5 mins, 5-95% B from 5-15 mins, 95% B from 15-20 mins, 95-5% B from 20-22 mins, and 5% B from 22-30 mins. The dominant transitions for each species are plotted as in Fig. 1.

**Supplemental Figure S2.**
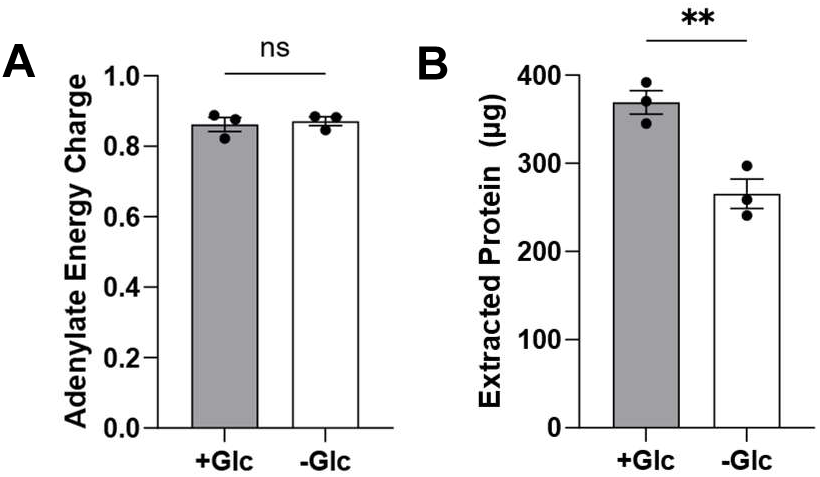
Adenylate energy charge (AEC) and total protein levels in glucose-starved HCT116 cells. HCT116 cells were cultured overnight with (*+Glc*) or without (*-Glc*) glucose and then analyzed for ATP, ADP, AMP. The adenylate energy charge (**A**) was calculated as AEC = (ATP + 0.5 * ADP) / (ATP + ADP + AMP). After metabolite extraction, the insoluble pellet was resolubilized for protein quantification. Mean ± sem, n = 3. ** p < 0.01 by unpaired t-test.

